# Cooler: scalable storage for Hi-C data and other genomically-labeled arrays

**DOI:** 10.1101/557660

**Authors:** Nezar Abdennur, Leonid Mirny

## Abstract

Most existing coverage-based (epi)genomic datasets are one-dimensional, but newer technologies probing interactions (physical, genetic, etc.) produce quantitative maps with two-dimensional genomic coordinate systems. Storage and computational costs mount sharply with data resolution when such maps are stored in dense form. Hence, there is a pressing need to develop data storage strategies that handle the full range of useful resolutions in multidimensional genomic datasets by taking advantage of their sparse nature, while supporting efficient compression and providing fast random access to facilitate development of scalable algorithms for data analysis. We developed a file format called cooler, based on a sparse data model, that can support genomically-labeled matrices at any resolution. It has the flexibility to accommodate various descriptions of the data axes (genomic coordinates, tracks and bin annotations), resolutions, data density patterns, and metadata. Cooler is based on HDF5 and is supported by a Python library and command line suite to create, read, inspect and manipulate cooler data collections. The format has been adopted as a standard by the NIH 4D Nucleome Consortium. Cooler is cross-platform, BSD-licensed, and can be installed from the Python Package Index or the bioconda repository. The source code is maintained on Github at https://github.com/mirnylab/cooler

## Introduction

Recent years have seen a ramp in production of large datasets that map associations between genomic loci. Of note are high-throughput chromosome conformation capture (3C) technologies (1, 2), such as Hi-C (3) and its variants, which produce two dimensional maps of chromosomal contacts. These technologies have undergone incremental improvements in technological resolution (cutting frequency, capture radius), biological sampling (cell numbers, library complexity) and technical sampling (sequencing depth), making it possible to resolve features at increasingly finer scales. Hi-C and related experiments also span a growing range of experimental scales, e.g. from single cells to large cell populations, unbiased vs. specific enrichment methods; for a review, see (4). As a result, there is a need for data structures that are flexible enough to accommodate data of massive size and varying degrees and patterns of sparsity, and easily adapt to new experimental techniques and novel metadata.

In the case of 3C-based experiments, pairs of sequence tags identify chimeric ligation junctions between DNA fragments. It is natural to subject these paired tags to *binning*, either by assigning them to putative restriction fragments, or more commonly, by aggregating them with respect to genomic intervals of some fixed size. Such gridded binning also suppresses count noise and increases effective coverage (5). The result is a quantitative genomic *matrix*, whose dimensional axes comprise a series of fixed or variable-length genomic intervals.

Today, processed Hi-C data and similar two-dimensional datasets are often still persisted using flat text files. For large and high-resolution datasets, this poses bottleneck challenges for basic processing, analysis and visualization. There exist compression and indexing strategies for tabular text files that mitigate these challenges to some degree by enabling random access (6). However, binary formats can provide more efficient and compressible storage, faster I/O, and preserve numerical precision. Several custom binary formats have been developed for Hi-C data, including butlr (7), hic (8), and MRH (9). They are useful in that they organize the data more efficiently and permit random access, but their strict byte layouts makes them rather inflexible for accommodating different data types, metadata or additional information. A popular alternative is the HDF5 container format (10), which provides the freedom to organize collections of editable multidimensional array data and metadata in binary form in a hierarchy similar to that of a file system. HDF5 is referred to as “self-describing” because objects can be inspected for their storage metadata, such as type, compression, and array shape. This enables it to serve as a flexible container for specific applications without constraining users to a strict preordained data organization. These features and its performance have made HDF5 very popular for storing large scientific datasets, and it has been made its mark in the Hi-C field for some time, in software packages such as hiclib (11), hifive (9), gcMapExplorer (12), HiCExplorer (13) and cworld (https://github.com/dekkerlab/cworld-dekker).

All the HDF5-based Hi-C formats in the tools mentioned use dense representations, i.e. full two-dimensional arrays of counts or transformed counts – including zeros for unobserved interactions; however, this strategy scales poorly with finer binning, whereby both size and sparsity of the data increase. For example, if we sample one billion contacts between kilobase-long fragments spanning the human reference genome from a population of cells, we are guaranteed to fill less than 1% of the trillions of elements in the available 2D space. Moreover, DNA contact frequency also exhibits a characteristic density pattern whereby contacts are much more densely sampled near the diagonal in *cis*. Sparse representations are not only critical for scalable storage: algorithms such as matrix balancing and PCA can be adapted to operate using only non-zero elements, and very finely binned maps can be used to look at patterns averaged over many genomic loci. Storage that accommodates a wide range of data resolutions is also necessary to visually explore the full depth of scale of such large datasets.

Here we present a data model, an implementation and a support library for a sparse, scalable HDF5-based genomic array format called cooler. We first describe a general sparse data model for genomically-labeled arrays, designed with Hi-C data in mind, but flexible enough to accommodate any binned genomic data describing associations, correlations, or interactions, such as linkage disequilibrium statistics. We then present its implementation as a file format in HDF5 that can support genomic matrices at any resolution as well as multi-resolution files, with a support software library in Python, also called cooler. The library provides the functionality to create, aggregate and manipulate the contents of cooler files, provides an application programming interface (API) to materialize genomic range queries in both tabular and array forms. Its design supports both sequential and random access, ideal for the development of out-of-core data processing algorithms. A command line interface is shipped with the cooler package for convenient scripting, application and pipeline integration.

## Data model

We outline a simple but flexible data model for representing multidimensional binned genomic data termed genomically-labeled sparse arrays (GLSAs). In the context of data structures, matrices and tensors are normally referred to as *arrays*, their individual shape dimensions are sometimes also termed *axes*. We will refer to the discrete units along the array axes as *bins*. Throughout we assume that the array axes are *homogeneous*, i.e. correspond to the same genome assembly, binned in the same way. A two-dimensional genomically-labeled array is a data structure that assigns unique quantitative values to pairs of bins obtained from an interval partition of a reference genome assembly. These pairs of bins make up the *coordinates* of the array’s elements. Because they are often visualized as heatmaps, we also use the term “pixels” to refer to elements of 2D arrays. By omitting pixels possessing zero or no value, the representation becomes sparse.

A genomically-labeled array of dimension two or greater can be represented with a single table similar to the BEDPE format, where each non-zero array element is described by a record listing the genomic coordinates of its bins and additional bin-related attributes alongside the element’s quantitative value(s) (Figure 1b, right). In Hi-C, for example, additional bin-related quantities include normalization weights or A/B compartment signal. However, at the scale of Hi-C data, this representation is limited by the fact that bin-related attributes (coordinates, weights, etc.) can be duplicated many times throughout the table, in numbers greatly exceeding the total number of bins.

**Fig. 1.**
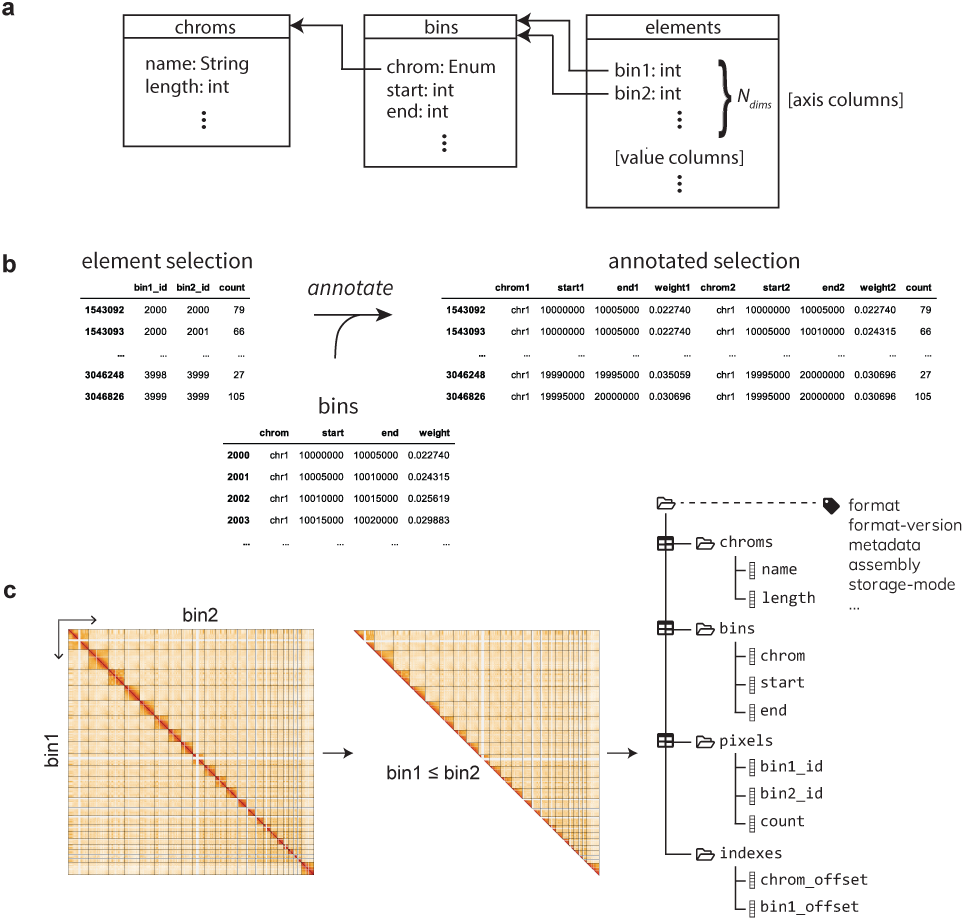
Data model for genomically-labeled sparse arrays and cooler format. **a,** Diagram of the GLSA data model. A multidimensional genomically labeled array can be represented via a decomposition that distinguishes the attributes describing the genomic intervals (table labeled *bins*) that make up the coordinates of the array’s axes from the actual non-zero elements of the array (table labeled *elements*). The element table contains one or more numerical *value columns* and simple integer coordinates that reference rows of the bin table (depicted using arrows). The bin table’s records describe a sequence of ordered, non-overlapping genomic intervals, minimally described by the reference sequence (*chrom*), and *start*, and *end* positions. The *chrom* column is further encoded as an integer enumeration to reference a third table labeled *chroms*, which contains attributes describing the reference sequences themselves, such as their genomic lengths. **b,** Any selection of rows of the element table can be annotated by joining with the appropriate columns of the bin table. **c,** For symmetric matrices, such as Hi-C maps, only upper triangular pixels are stored to eliminate duplication. Right, a diagram of a cooler data collection’s hierarchical structure. The three tables are modeled as HDF5 groups (depicted as folders) while the table columns are stored as 1D arrays, which are chunked and compressed internally by HDF5. A reserved set of metadata HDF5 attributes are associated with the root group of the data collection, including a flag indicating whether the matrix is to be interpreted as symmetric.

We eliminate such redundancy by replacing the single table with two separate ones (Figure 1a). The first is a *bin table* that describes the complete genomic bin segmentation on both axes of the matrix, such that each bin-related attribute is fully described by a column, or one-dimensional array. The second table contains distinct axis columns (`bin1_id`, `bin2_id`) that reference the rows of the bin table: the resulting *pixel table* is a condensed representation of the nonzero elements of the array. Conveniently, this corresponds to the classic coordinate list (COO) representation for sparse arrays (14). For completeness, we include a third *chromosome table* to list the chromosomes (or other scaffolds) of the genome assembly, their genomic lengths, and any other chromosome-specific attributes. Note that while the bin and chromosome tables tend to have a relatively small memory footprint, it is the pixel table that can greatly exceed memory storage for large datasets.

Furthermore, for symmetric matrices, such as Hi-C, we reduce data requirements and preserve a unique representation by keeping only unique pixels and orienting them to lie within the upper triangle of the matrix, discarding their lower triangular transpose elements (Figure 1c). Altogether, for a given ordering of the chromosomes of an assembly, this decomposition uniquely represents a given genomically-labeled 2D array.

Given any collection of rows from the pixel table, its full representation (or any subset of bin-related attributes) can be recovered inexpensively at the application level by performing relational joins between the axis columns of the pixel table and the desired bin attributes in the bin table. We refer to this as element *annotation* (Figure 1b).

## Cooler format

We implemented the GLSA data model for 2D arrays using HDF5, defining a file format called cooler, with the recom-mended extension `.cool`. A full specification can be found in the online documentation. We outline the key elements in this section.

### Specification

HDF5 files are organized hierarchically, akin to a file system within a file, with two primary structures, *groups* and *datasets*. Groups serve the role of directories and contain zero or more groups or datasets. Datasets are multi-dimensional arrays, which can be flexibly sized, chunked and passed through various I/O filters, including checksumming and compression. Both groups and datasets can be assigned key-value user metadata, called *attributes*, and like POSIX file systems can be referenced with relative or absolute paths separated by slashes.

HDF5 does not natively support sparse arrays or relational data structures: its datasets are dense multidimensional arrays. Therefore, we model a table as a group of equal-length 1D arrays representing columns. Although HDF5 does support row-oriented storage using compound data types (i.e. structured arrays), we chose to implement column-oriented storage because of the advantages it provides, including cheap addition or removal of columns, efficient slicing along columns, and more efficient compression (15). Our simple table model thus allows for easy addition of columns and appending of rows, but not random insertion of rows. This was deemed reasonable since the raw datasets are normally write-once, but global data transformations are not uncommon. Moreover, it does not enforce a specific ordering on the columns, though a conventional order for required columns is provided in the specification.

A schematic of the array hierarchy representing a single matrix is shown in Figure 1c. Three HDF5 groups representing the *chrom, bin* and *pixel* tables live directly beneath the collection’s root group. Our specification permits any number of additional columns in the three tables and any additional group hierarchies and metadata, as long as they are not nested within the group of a table. For example, additional 1D signal tracks can be appended to the bin table, and pixel tables can contain multiple value columns (e.g., the color bands of an image). An additional group called *indexes* lives alongside the three tables and contains two index arrays described below.

We term the complete object hierarchy representing a matrix a *data collection*. The specification requires that some standard metadata be provided in the attributes of the data collection’s root group (Figure 1c). We reserve an additional attribute slot for storing serialized custom user metadata in JSON format: for example, experimental details, processing logs or mapping statistics. Since all versions of the HDF5 library ship with zlib compression, for maximum portability, it was chosen as the default compression filter for all columns.

### Indexing

We further stipulate that the records of the pixel table must be sorted lexicographically by the bin ID along the first axis, then by the bin ID along the second axis. This way, the *bin1_id* column can be substituted with an array of offsets that serves as a lookup index for the rows of the matrix, stored under *indexes/bin1_offset* (Figure 1c). With this index, we obtain a compressed sparse row (CSR) sparse matrix representation (14). Given an enumeration of chromosomes, the bin table must also be lexicographically sorted by chromosome then by start coordinate. Then similarly, the *chrom* column of the bin table will reference the rows of the chrom table, and can also be substituted with an offset array, stored under *indexes/chrom_offset* (Figure 1c). Because of the efficient compression of sorted columns, we preserve the original *bin1_id* and *chrom* columns so that the indexes may be dispensable for a reader that does not wish to use them.

### Flavors

Although the standard cooler file contains a single data collection located at the file’s root group, we allow a cooler file to store multiple data collections anywhere in the group hierarchy of an HDF5 file, as long as they are not sequentially nested. Any number of data collections is permitted, and each individual data collection can be referenced using its qualified group path. We therefore support a flexible URI syntax to locate data collections within a file (see below) and provide two conventional “flavors” of the file layout: the single resolution and multi-resolution or “zoomified” cooler (often suffixed `.mcool`), which is ideal for interactive multiscale visualization, as exemplified by HiGlass (16) (Figure 2).

**Fig. 2.**
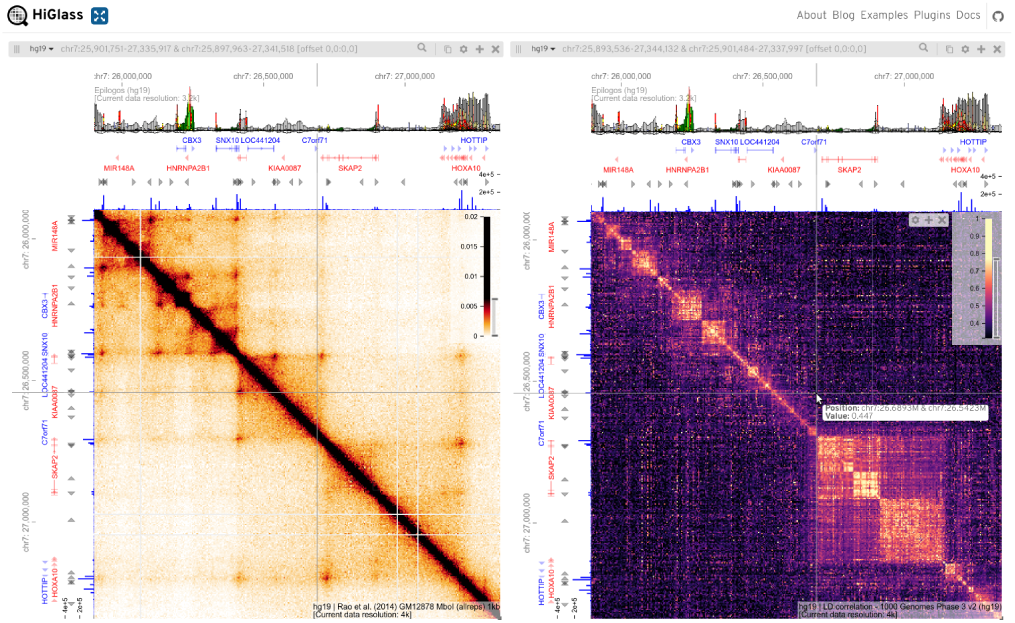
Multiresolution cooler files can be synchronously browsed with continuous pan and zoom. Snapshot of the HiGlass web application (16) displaying two zoomified (multiresolution) cooler files storing different 2D genomic data types in two “views” linked by location and zoom level with synchronized panning and zoom. HiGlass provides a continuous genome-wide Google Maps-like user interface that automatically and smoothly adjusts data resolution. The Hi-Glass client requests the necessary data to render heatmap tiles from a Python-based server using the cooler library. The views contain additional genomic and epigenomic tracks along their top and left borders, also synchronized. Left: a Hi-C map for GM12878 lymphoblastoid cells from (17). Right: a heatmap dis-playing locally averaged linkage disequilibrium (LD) *r* values between all pairs of human variants derived from the 1000 Genomes Project, Phase 3 (18). Pair-wise LD statistics were calculated genome-wide on all variants over a 5 Mb sliding window using tomahawk (https://github.com/mklarqvist/tomahawk) and the output was ingested into a cooler file using `cooler cload` at a 1 kb-resolution aggregation and further aggregated using `cooler zoomify`. Borders: hg19 coordinates and gene annotation tracks as well as CTCF ChIP-seq bigwig signal tracks (blue bars) from ENCODE (19). The top border track displays Epilogos (https://epilogos.altiusinstitute.org/) tracks summarizing ChromHMM state annotations over 127 cell types in the Roadmap Epigenomics Project’s Core 15-state model (20). This figure can be browsed interactively online at https://higlass.io/app?config=W4DNgqjXRNWPQ7Nbz7NLnQ.

## Cooler package

We provide a Python-based convenience library to manage cooler data collections. It provides tools to create and append data to collections, to merge, aggregate and normalize them, and to and query their contents and metadata. To identify cooler data collections that may lie at any level of the group hierarchy of a file, we support a URI syntax consisting of a file path, optionally followed by double colon ::followed by a fully qualified HDF5 group path. If the double colon and group path are omitted from the URI, the data collection is interpreted as being located in the file’s root group.

### Command line interface

The cooler Python package ships with a command line interface (CLI) for most common manipulations (Figure 3a). They are provided as subcommands under a top-level `cooler`command namespace. We briefly describe the main commands below, grouped by theme. All these actions are also available and further customizable pro-grammatically through the Python API.

**Fig. 3.**
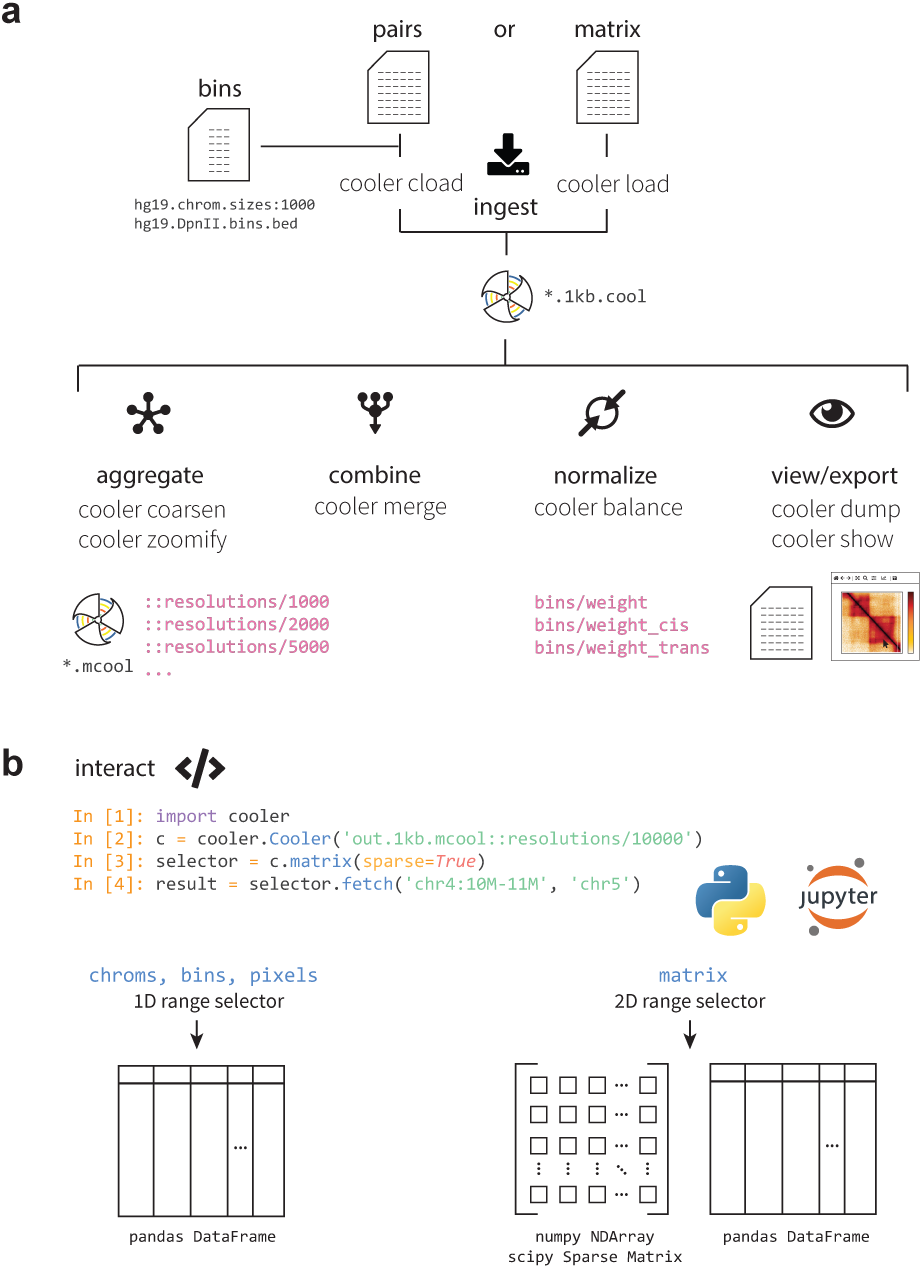
Cooler command line interface and Python library. **a,** Summary of the main categories of `cooler`commands available with the cooler Python package, illustrating the flow of data. The main operations include the ingestion of file or text streams to create new coolers, aggregation and coarsening of existing coolers to lower resolutions, merging of axis-compatible matrices, normalization of cooler matrices by iterative correction, utilities to serialize and stream out the data and metadata inside a cooler file and to process range queries, and a lightweight viewer to visually inspect a matrix. For example, one uses either the `load`command to ingest pre-aggregated data already in matrix form, or the `cload`command to aggregate paired tag records into a matrix. The genomic bin segmentation defining the axes of the matrix must be provided separately by providing either a path to a BED file or a path to a chromosome sizes file along with a specified fixed bin size. **b,** The cooler Python library provides a `Cooler`class that exposes data *range selectors* to facilitate data retrieval and analysis. The individual *chrom, bin* and *pixel* tables are accessible using 1D range selectors that accept column and row-range selections and yield pandas data frame output. A cooler’s matrix values are also exposed using a 2D range selector that processes range queries provided either as genomic coordinate intervals in UCSC-style notation (using the `fetch`method) or as integer matrix coordinates (using Python slice syntax). The retrieved 2D range data may be materialized as dense NumPy arrays, sparse matrices, or data frames. For symmetric coolers, the file’s upper triangular data will be appropriately mirrored in the array and sparse matrix outputs.

### Ingestion

There are several commands used to create cooler data collections from input data. The first required input is a definition of the bin segmentation of the genome assembly to which the interaction data was mapped. This can be provided either as a BED file, or in the case of a fixed bin size resolution, alternatively as a file listing the chromosome lengths (a chrom-sizes file) along with the bin size (resolution) in base pairs. The second required input is either (1) a list of records describing aggregated data (i.e. a matrix) or (2) a list of un-aggregated paired tags.

If the input data are unaggregated paired tags, the command `cooler cload`will aggregate the pairs according to the provided bin segmentation. If the input is already binned, the `cooler load`command will convert the input matrix into the cooler format.

### Aggregation

Cooler data collections at one or more base resolutions can be aggregated or *coarsened* to lower resolutions using the `cooler coarsen`command. Moreover, recursively aggregated multi-resolution cooler files can be generated using the `cooler zoomify`command. The resulting files are suitable both for data analysis and for multiscale visualization (21).

### Merging

Any combination of cooler data collections with compatible bin axes, such as technical or biological replicates of an experiment, can be pooled together in a memory-efficient manner using the `cooler merge`command.

### Balancing

The *de facto* standard method for normalizing Hi-C data is matrix balancing, also known as iterative correction (11). Because is it such a common transformation, we include this functionality in the cooler package. The output is a vector of balancing weights (the reciprocal of the biases described in (11)). These weights are applied by multiplying each pixel value by the weights associated with its two genomic bins.

The `cooler balance`command performs bin-level filtering and balancing using a parallel and out-of-core implementation of iterative correction with extensive options. Balancing weights are normally stored as columns in the bin table of a data collection and applied on-the-fly during querying. Alternatively, one can generate a new data collection of transformed counts.

### View/Export

The contents of the *chrom, bin* and *pixel* tables may be serialized as delimited text using the `cooler dump`command, which also supports genomic range queries and pixel annotation. Additionally, the `cooler show`command provides a lightweight interactive matplotlib visualization to inspect and explore the data.

### Python library

The cooler library provides programmatic access to all of the above functions as well as an API to select arbitrary ranges of data from the tables and perform 2D range queries on matrices (Figure 3b), powered by the *h5py* (22) Python interface to HDF5. It provides both tabular and 2D array-based interfaces to the sparse (and, for symmetric matrices, upper triangular) data representation in the file. It understands both global array indices and genomic coordinate ranges.

To integrate seamlessly into the Python data ecosystem, the cooler package’s API materializes queries in several common scientific data structures, including dense *NumPy* arrays, *SciPy* sparse matrices and *Pandas* data frames. Furthermore, selections of pixels in data frame form can be annotated with genomic bin columns as depicted in Figure 1b using the *an-notate* function.

The combined CLI and API of the cooler package makes it suitable for use in scripts and pipelines and for inclusion in other software libraries. Thanks to its ease of integration with Python-backed tools, it has already been successfully integrated into 3D genome browsers, pipelines and visualization tools, including HiGlass (16), the WashU Epigenome Browser (23), HiCExplorer (13), and HiCPlotter (24). The cooler package also provides the flexibility to facilitate interactive data analysis in environments such as the Jupyter Notebook platform (25). A comprehensive suite of tools dedicated to Hi-C data analysis using cooler files is being developed as part of a package called *cooltools* and will be presented else-where.

### Implementation

Cooler is implemented as a Python package and supports Python versions 2.7 and 3.4 or greater and works on Linux, Mac OS X and Windows platforms. The cooler package is open source, BSD licensed, and the source code is maintained on GitHub at https://github.com/mirnylab/cooler. The documentation is hosted at https://cooler.readthedocs.io.

Cooler can be installed using Python’s pip package manager,

`$ pip install cooler`

or from the bioconda distribution channel (26) using the conda package manager.

`$ conda install -c bioconda cooler`

For docker users, the command line interface is also available through BioContainers (27).

$ docker run quay.io/biocontainers/cooler cooler –help

## Discussion

We present a sparse data model and file format for genomically-labelled arrays with minimal redundancy but enough flexibility to support a wide range of data types, data sizes and future metadata requirements. A sparse representation in particular is crucial for developing robust tools and algorithms for use on increasingly high-resolution multidimensional genomic data sets that need to operate on subsets of data at a time. While we developed a command line suite and Python API to create, inspect and manipulate cooler data collections, we also set out to define a simple enough specification that it could be easily interpreted across programming environments using the established APIs of the underlying storage layer.

In selecting a storage layer on which to implement our sparse array data model, HDF5 was chosen because it is an open-source, portable and performant format for scientific data with widespread use and whose self-describing generic data structures are well suited to data modeling. Furthermore, we required exchangeable encapsulated files, rather than sharded files or distributed databases, because the former are still indispensable for modern bioinformatic workflows and data analysis practices. Popular binary formats such as the current versions of MATLAB’s mat files and Unidata’s NetCDF4 are built as abstractions on top of HDF5 (28) and it has been used to store petabytes of mission critical data, such as NASA’s Earth Observation System, for decades (29). Importantly, HDF5 supports chunking and compression, arrays of unlimited size, efficient array subset selection (slicing), and high level APIs exist for a wide variety of programming languages, including C/C++, Java, Python, Perl, and R.

Nevertheless, the high-level GLSA data model can be implemented using a variety of storage strategies for different goals. For example, because the model minimally fragments the data while eliminating duplication, it can be very useful for text-based interchange for multidimensional genomic arrays or parts thereof. Indeed, for Hi-C data, a two-file text format based on a bin file and upper triangular element file was introduced by the HiCPro pipeline (30). Other open source backends in which GLSAs could be implemented include Apache Parquet (https://parquet.apache.org/), a cloud-optimized columnar binary format for big tabular data; an emerging array storage technology called TileDB that provides native high performance sparse array support (https://tiledb.io/); and a new format called Zarr (https://github.com/zarr-developers/zarr) that provides an HDF5-inspired implementation of chunked, compressed, N-dimensional arrays that can work on top of a variety of storage layers (file system hierarchies, zip files, or key-values stores). Such new technologies are poised to become more important as community genomic data migrates increasingly to cloud storage and computing environments.

Of course, our design decisions involved certain tradeoffs. The data model assigns every genomic bin a global bin ID, which is sensitive to the chromosome order chosen. This tradeoff was deemed preferable to more complicated schemes that add additional bin identifiers or that divide each scaffold-pair block into a separate group. HDF5 supports a row-based storage model for tables using compound data types (structured arrays), but the compression, append, and performance benefits of columnar were deemed important enough to define a column-based storage model. The CSR indexing scheme in our HDF5 schema is very space-efficient, but not optimal for 2D range queries because the data are not serialized in a way that strongly preserves 2D locality. However, by using different sort orders on the element data, the data model we present can support more sophisticated indexing schemes, such as space-filling curves (31). Finally, the data model implemented describes two-dimensional arrays with homogeneous sets of axes. Though not yet implemented in the cooler package, handling heterogeneous axes is a matter of including separate chromosome and bin tables for each distinct axis. The data model also extends naturally to multi-dimensional tensors by including additional axis columns in the element table.

Cooler provides a scalable solution to tackling the analysis and visualization of big Hi-C data and other genomically-labeled matrices at any data resolution, scaling to massive data sizes without incurring the read and write bottlenecks of dense storage. It is also amenable to external-memory (often called out-of-core) algorithms that can be controlled to maximally exploit multiple cores and disk I/O while not overwhelming memory resources (32). The cooler package’s command line tools facilitate integration into scripts and workflows, and its Python API allows users to leverage the powerful resources of the Python data ecosystem while the portable HDF5 format makes it readily accessible in other environments such as Java, and R. The cooler format has adopted as a standard for Hi-C data storage, along with hic (8) and the pairs text format (https://github.com/4dn-dcic/pairix/blob/master/pairs_format_specification.md), by the 4D Nucleome Consortium’s Data Coordination and Integration Center, and is already supported by a number of genomic analysis and visualization tools, including HiGlass (16).

## ACKNOWLEDGEMENTS

We greatly thank Anton Goloborodko and Maxim Imakaev for many extensive discussions, contributions to code development, and providing feedback on the manuscript. We are grateful to Peter Kerpedjiev for helpful discussions and for Hi-Glass integration. We also thank Geoffrey Fudenberg, Sergey Venev, Ilya Flyamer, Joachim Wolff, members of the Mirny lab, and members of Nils Gehlenborg’s and Peter Park’s groups for feedback, contributions, and animated participation on the public issue tracker. We acknowledge support from the National Institutes of Health Common Fund 4D Nucleome Program (DK107980), NIH (GM114190), and the National Science Foundation (Physics of Living Systems: 1504942). This preprint is formatted using a LATEXclass by Ricardo Henriques (CC-BY).

